# The variations of human miRNAs and Ising like base pairing models

**DOI:** 10.1101/319301

**Authors:** Jyoti Prasad Banerjee, Jayanta Kumar Das, Pabitra Pal Choudhury, Sayak Mukherjee, Sk. Sarif Hassan, Pallab Basu

## Abstract

miRNAs are small about 22-base pair long, RNA molecules are of extreme biological importance. Like other longer RNA molecules, messages in miRNAs are encoded by the permutations of only four nucleotide bases represented by A, U, C and G. However, just like words in any language, not all combination of these alphabets make a meaningful word. In fact, we find that the distributions of nucleotides bases in human miRNAs show significant deviation from randomness. First, a miRNA sequence containing four bases are mapped into a binary string with three kinds of classifications according to their chemical properties. Then, we propose a simple nearest neighbor model (Ising model) to understand the statistical variations in human miRNAs.

## 1. Introduction

Micro-RNAs (miRNAs) are small non-coding RNA molecules. They are made of the different permutations of nucleotide bases A, U, G and C. The number of bases in the miRNA is conserved across the species and they vary around 22 nucleotides (mostly). They play important role as the regulators of gene expression and they target some of messenger-RNAs (mRNAs) [1, 2, 3]. Mainly, the function of miRNAs is to down-regulate gene expression [4]. In recent years, the effects of miRNAs are also found in malignant cells; the miRNAs influence numerous cancer-relevant processes in a malignant cell [5, 6]. It will be worthy if the governing role of miRNAs could be apprehended from nucleotide bases and their properties. The recent study have demonenstrated how the miRNAs of five species are distributed on the basis of purine-pyrimidine bases that used different mathematica parameters [7]. To have a deeper understanding of the nature of miRNAs, here we study a few statistical properties of the miRNAs and propose some simple physics-inspired models to account for their ensemble properties. As we will discuss, our results may shade some light on how the miRNAs are produced in the cell.

## 2. Results and Discussions

### 2.1. Dataset specification

The whole miRNA sequences (for *Human*, Hominidae family) are accessable through the website (a miRNA database: http://www.mirbase.org/). In human, 2588 reported mature miRNAs sequences are found. Their lengths vary from 16 to 28 nucleotide bases, but mostly within the range of 20-24 bases (Fig 1). Let *L* be the data set containing 2588 miRNAs i.e. *L* = 2588. The average length or mean (*M*(*L*)) of 2588 miRNAs is 21.5881. All the computational codes are written in *MATLAB R2016a* software.

**Figure 1:**
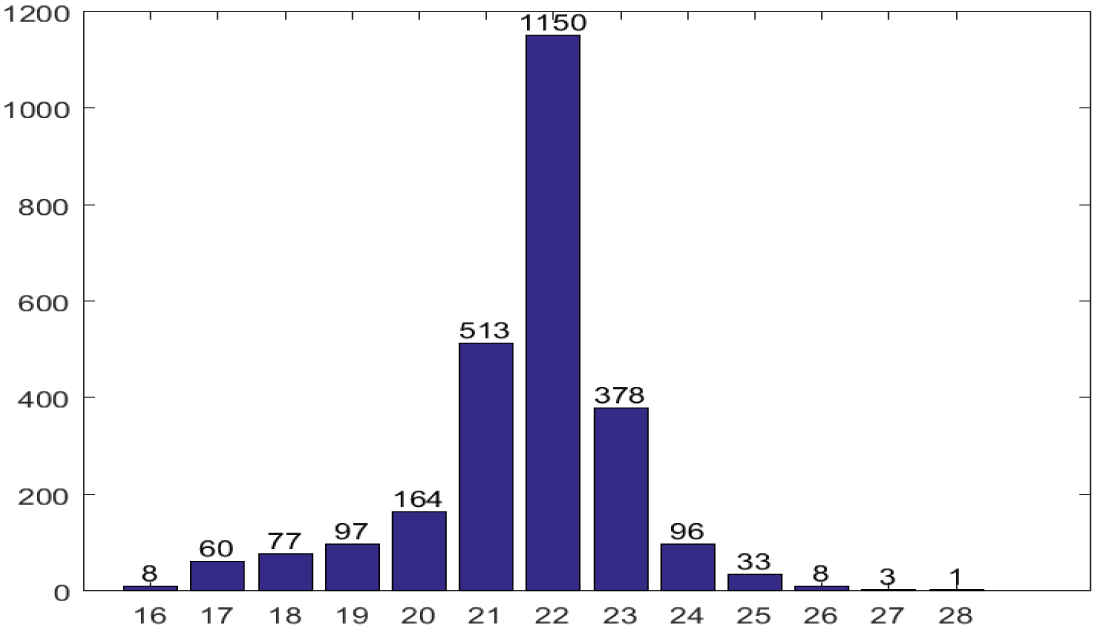
The length variation and the respective number of miRNAs count from the total of 2588 miRNAs.

### 2.2. Individual variation of nucleotides bases

There are only four ribonucleotide bases which are distributed over the miRNAs. Here, we will see their distribution in terms of the probability measure.

Let, *Pr*(*X*) is the probability of a nucleotide *X* (*X* ∈ {*A, G, U, C*}) to be present in the 2588 miRNAs, then

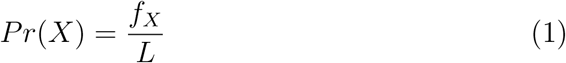

where *f_X_* is the average number of occurrence of the nucleotide *X* over 2588 miRNAs.

The average frequency distribution of all 2588 miRNAs are shown in Fig 2.

**Figure 2:**
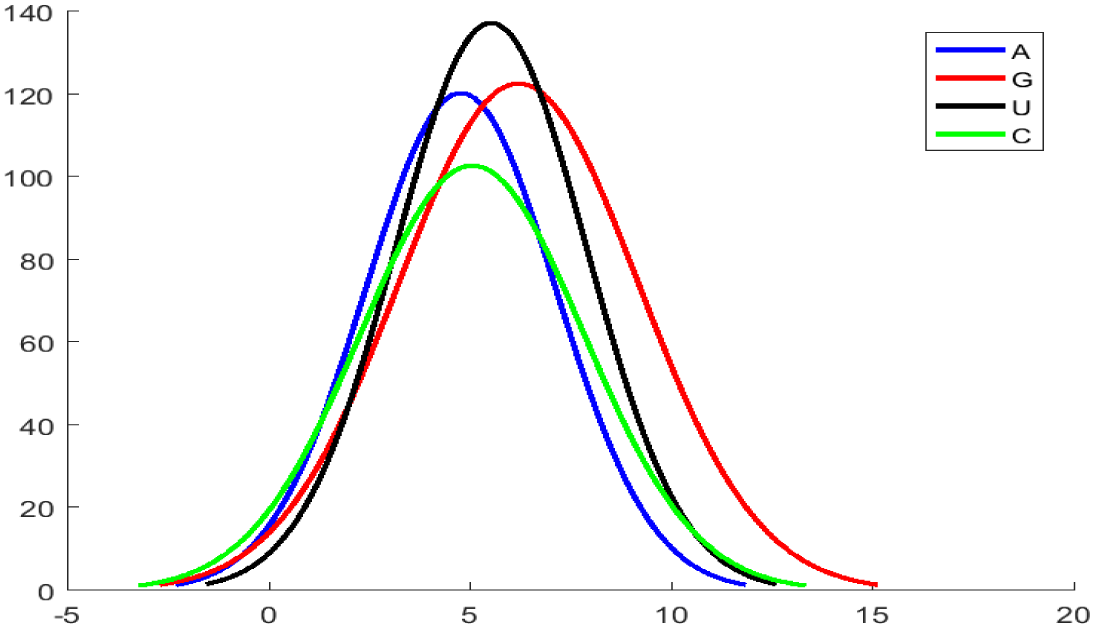
The average frequency distribution of 4 nucleotides over 2588 miRNAs.

From the data set, we find *f_A_* = 4.77, *f_G_* = 6.21, *f_U_* = 5.55 and *f_C_* = 5.53. So, *Pr*(*A*) = 0.22, *Pr*(*G*) = 0.28, *Pr*(*U*) = 0.26 and *Pr*(*C*) = 0.24. It is observed that *Pr*(*A*) ≠ *Pr*(*G*) ≠ *Pr*(*U*) ≠ *Pr*(*C*). Therefore, we find the individual variations of each nucleotide base over the 2588 miRNAs. And the orderis *Pr*(*A*) < *Pr*(*C*) < *Pr*(*U*) < *Pr*(*G*).

### 2.3. Grouping of nucleotide bases and transformations into binary

The four-letter alphabets (A/G/U/C) may be mapped onto two-letter (binary) alphabets in three distinct possible ways. As it turns out, these three classifications may be understood on the basis of chemical properties of nucleotide bases [8]. These are:

- **Purine-Pyrimidine classification:** The bases AG and UC are the purine and pyrimidine (Pu-Py) groups respectively.
- **Strong-Week H-bond classification:** The bases CG and AU are the strong H-bond and week H-bond (St-We) groups respectively.
- **Amino-Keto classification:** The bases AC and GU are the amino and keto (Am-Ke) groups respectively.

Among the three classifications, the purine-pyrimidine class is possibly the most important one [9]. Purines are the most widely occurring nitrogen-containing heterocyclic compounds in nature. In order to form DNA and RNA, both purines and pyrimidines are needed by the cell in approximately equal quantities. But, this may not hold true for miRNAs.

For all of these three groupings, we can assign the bits 1’s and 0’s to the respective members of any of the aforementioned classifications. For example, for the purine-pyrimidine class we may have the following rule:

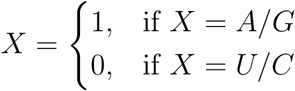

The mean occurrence for all the different classification From the dataset, for purine-pyrimidine grouping *M*(*Pu*) = 10.99 and *M*(*Py*) = 10.59, for strong-week H-bond grouping *M*(*St*) = 11.27 and *M*(*We*) = 10.31, for amino-keto grouping *M*(*Am*) = 9.84 and *M*(*Ke*) = 11.74. The occurrence distribution of Pu, St and Am is shown in Fig 3. The cumulative distribution function for these three groupings are plotted in Fig 4.

**Figure 3:**
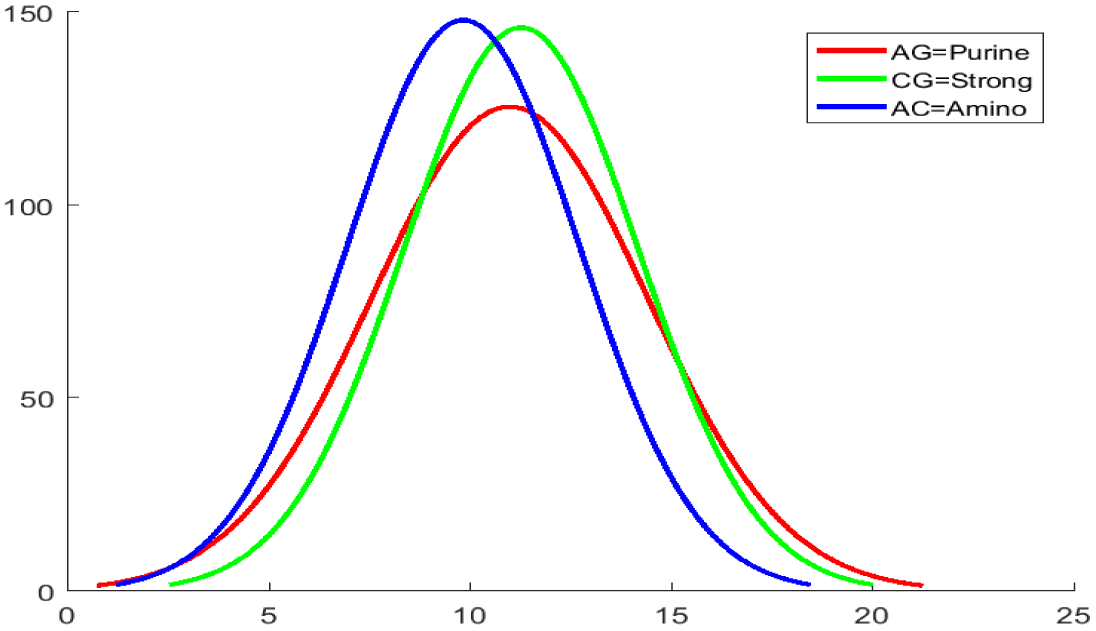
The occurrence distribution of Purine (AG), Strong (CG) and Amino (AC) classes.

**Figure 4:**
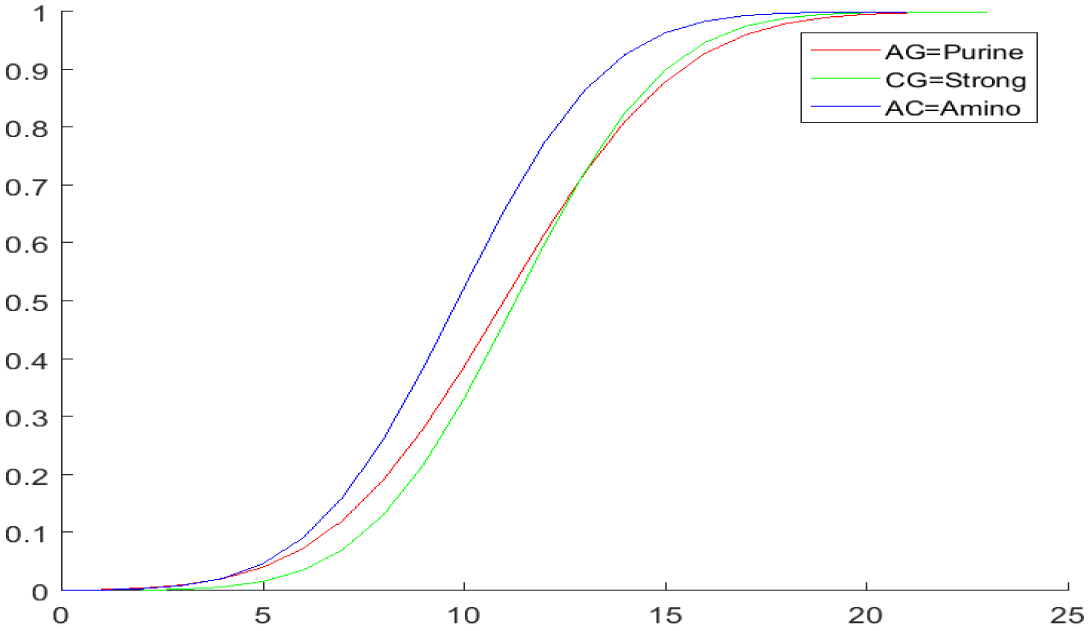
The cumulative distribution function (cdf) plot of Purine (AG), Strong (CG) and Amino (AC) classes.

Interestingly, for all three categories, the distribution of occurrence of 0 (or 1) in the binary string representation of miRNAs fits well to a normal distribution. The group-specific distributions of the nucleotide base pairs vary significantly from each other. However, we observe that the mean occurrence of any member (pu or py) in the Purine-Pyrimidine classification is the closest to the occurrence averaged over all the members of all the three classifications (10.79), as compared to other two groupings. The mean and variance for each members of all the classifications are provided in the Table 1. The question that naturally arises is: which kind of model best describes the observed variances in the distribution of nucleotide bases?

**Table 1:** Expected variance and observed variance for the bases of each classification.

| Class | n | p | q | Mean<br>(np) | Variance<br>(npq) | Observed<br>Variance |
| --- | --- | --- | --- | --- | --- | --- |
| Pu-Py | 21.59 | 0.509 | 0.491 | 10.988 | 5.40 | 11.7 |
| St-We | 21.59 | 0.522 | 0.478 | 11.268 | 5.39 | 8.48 |
| Am-Ke | 21.59 | 0.455 | 0.544 | 9.839 | 5.35 | 8.26 |

### 2.4. Binomial Model

The simplest thing to do is to fit a binomial model. In a binomial model, probability of occurrence of a 0 (or 1) at each location is determined by an independent coin-toss with a probability *p* (or *q*). Here *p* + *q* = 1. The mean of an binomial distribution is *np*. The binomial distribution has a simple property that variance is *npq* (Theoretical/Expected). Hence from the mean we can calculate *p* and check if the variance is close to the observed variance. (Here, *n* = *M*(*L*) = 21.5881). The details of Expected variance and observed variance for the bases of each classifications are shown in Table 1.

So, from the above results we find that the observed variance is greater than the variance obtained from the binomial model. However, the discrepancy in variance for the Pu-Py classification is much larger than that for the other classifications. For all other classes the expected variance is much closer to the observed variance. So, there are patterns of miRNAs where one group (Pu or Py) is more frequent than what is expected from chance alone. Therefore, it is interpreted that miRNAs have a stronger interdependency with respect to the purine-pyrimidine class.

To have a better statistical understanding of our claim that observed mean is different from the mean calculated from observed variance assuming binomial distribution, we performed one-sample *t*-test (*h*) with the null hypothesis that two observed mean are equal. We performed the t-test separately for all three cases and calculated the *p* value. In all three cases we can reject the null hypothesis with high significance (Table 2).

**Table 2:** Observed mean, Expected mean, *p* value of each class for different cases.

| Class | Cases | Observed<br>mean<br>($O_m$) | Expected<br>mean<br>($E_m$) | P- value | Om/Em |
| --- | --- | --- | --- | --- | --- |
| Pu-Py | 00c | 5.54 | 4.96 | 0.0024 | 1.12 |
|  | 01c | 4.55 | 5.14 | 2.86E-152 | 0.89 |
|  | 10c | 4.47 | 5.14 | 6.39E-173 | 0.87 |
|  | 11c | 6.02 | 5.33 | 5.66E-23 | 1.13 |
| St-We | 00c | 4.06 | 4.7 | 3.31E-27 | 0.86 |
|  | 01c | 5.75 | 5.14 | 1.20E-102 | 1.12 |
|  | 10c | 5.61 | 5.14 | 2.81E-64 | 1.09 |
|  | 11c | 5.71 | 5.61 | 6.51E-13 | 1.02 |
| Am-Ke | 00c | 6.48 | 6.12 | 3.23E-09 | 1.06 |
|  | 01c | 4.68 | 5.12 | 3.09E-67 | 0.91 |
|  | 10c | 4.76 | 5.12 | 3.27E-46 | 0.93 |
|  | 11c | 4.66 | 4.26 | 3.51E-13 | 1.09 |

### 2.5. Mean-field model

In the previous subsection we have observed that the variance of the data is more than what is expected of the independent, binomially distributed binary strings. This discrepancy indicates that there could be interactions present within the elements of a string. These interactions may have a mean-field nature where every element of a binary string interacts with every other element within the same string with equal strength (all-to-all connected topology). In the other possible case, there might be a strong nearest neighbor interaction instead of a mean-field one. To check which of these two descriptions are closer to the observations from the data, we calculate and compare the probabilities of finding 2-clusters (i.e., like-pairs) of Purines and Pyrimidines in the binary strings obtained from the data (shown in Table 4) and the same obtained by the bootstrap method. In the bootstrap method we shuffle the elements of each binary string multiple times keeping the numbers of Purines and Pyrimidines constant. We then calculate the probabilities of a 2-cluster in the shuffled strings and obtain a distribution of them. If the probabilities obtained from the data falls under the bootstrap distribution, preferably close to the pick of the distribution, then one can infer that shuffling does not change the cluster-probabilities, and a mean-field picture is enough to model the statistical properties of the miRNA sequences. The distributions and the values calculated from the data are shown in figures 5 and 6.

A statistically significant difference between the observed means and values obtained in the bootstrap analysis indicate that the purines and pyrimidines are not positioned in a random fashion in the miRNAs. Their special arrangements, therefore, suggest that they have a rather strong nearest neighbour interaction.

**Figure 5:**
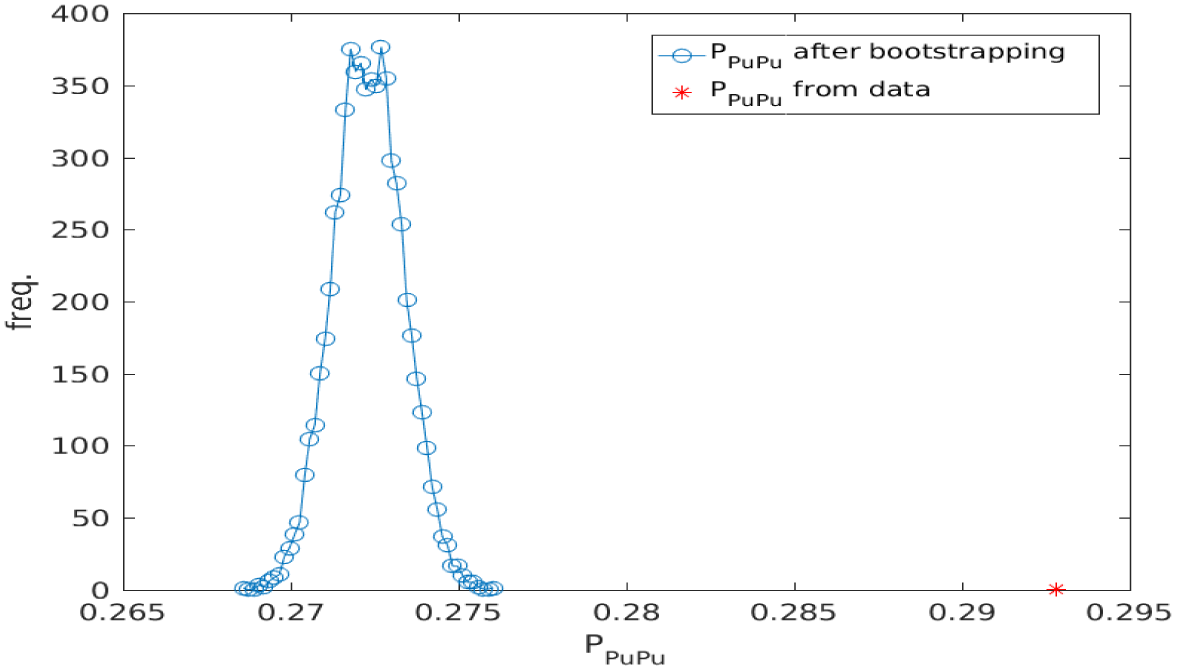
Distribution of probability of 2-clusters of Purines from bootstrap method and from the data.

**Figure 6:**
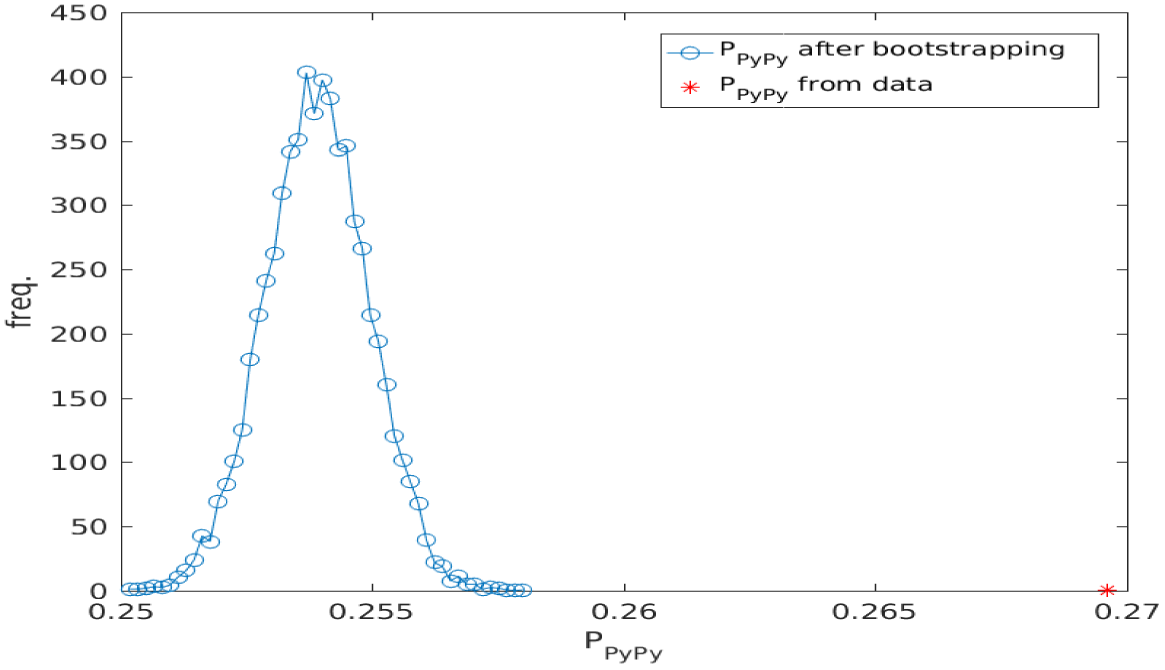
Distribution of probability of 2-clusters of Pyrimidines from bootstrap method and from the data.

**Figure 7:**
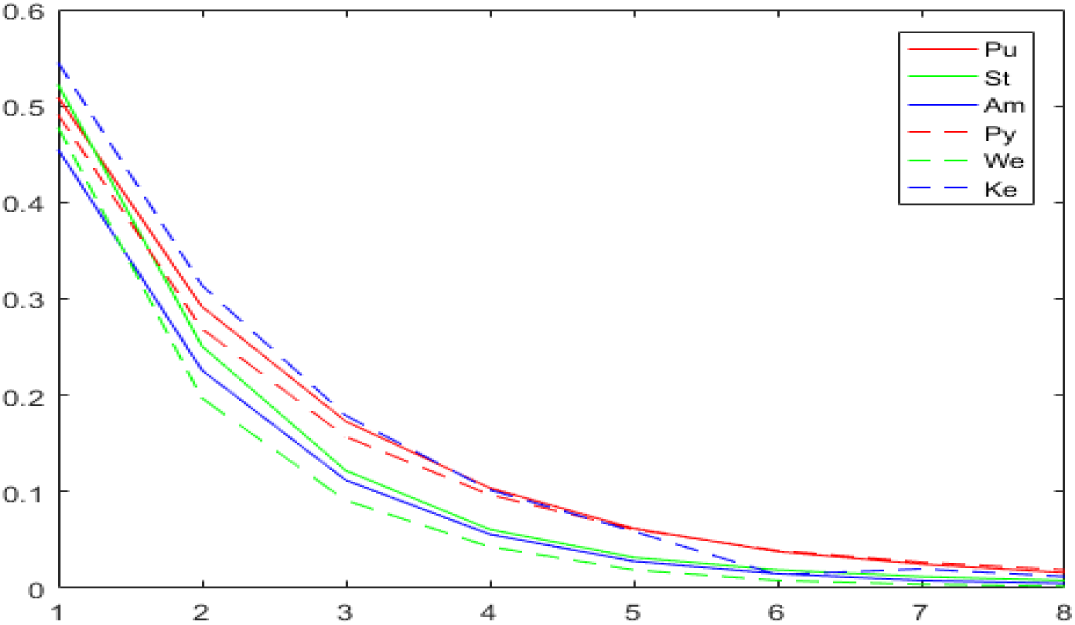
The probability distribution for the different lengths (1 to 8) strings of ones and zeros.

### 2.6. Nearest Neighbor Model: Ising Model

Here we construct a Nearest Neighbour (NN) model which assumes pair-wise interaction between any two nearest neighbour elements. In a different context, for various nucleotide sequencing data such models have previously been used [10]. In this model, the probability of occurrence of a particular string of length *N* with *r* zeros would be given by,

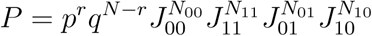

where ‘J’s are the strength of nearest neighbour interactions.

From Table 3 we observe that *P*_00_ ≈ *P*_11_ and *P*_01_ ≈ *P*_10_ for all 3 types of classifications. This indicates that it might be sufficient to club the four different interactions to interactions between similar elements and those between the different elements. Also, we know that *P*_0_ ≠ *P*_1_ (from Table 1) and *P*_00_ ≠ *P*_01_ (from Table 3) for all of the different classification schemes. These observations together suggest that a nearest neighbour model can possibly be a good candidate to fit and explain the observed data.

**Table 3:** Observed frequency and respective probability of each class for cases.

| Class | Cases | Frequency | Pr(00/01/10/11) |
| --- | --- | --- | --- |
| Pu-Py | 00c | 14341 | 0.27 |
|  | 01c | 11783 | 0.22 |
|  | 10c | 11575 | 0.22 |
|  | 11c | 15583 | 0.29 |
| St-We | 00c | 10497 | 0.20 |
|  | 01c | 14881 | 0.28 |
|  | 10c | 14524 | 0.27 |
|  | 11c | 13380 | 0.25 |
| Am-Ke | 00c | 16772 | 0.31 |
|  | 01c | 12112 | 0.23 |
|  | 10c | 12331 | 0.23 |
|  | 11c | 12067 | 0.23 |

In physics literature the nearest neighbour model is called the Ising model. Historically, the Ising model was created/used to describe the paramagnetic to ferromagnetic transition in 1-dimension. It is a semi-classical model which, in simple terms, assumes that the magnetic moments (spins) of atoms in a 1-datomistic chain can assume only 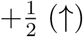 and 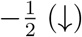 values in some suitable unit system along a particular (say, *Z*) axis. In a specific variant of the model, let us consider a closed (i.e., with periodic boundary condition) chain of N spins with nearest neighbour interaction strength, *J*, and chemical potential (magnetic field like term), *μ*. We assume that the inverse temperature, *β* = 1.

There can be two approaches to fit this model to the observed microRNA data. One is to calculate the susceptibility using the transfer matrix method and treat it as the variance of the difference between population of the two states (↑ and ↓ for Ising model, Pu and Py for our classification, for example). This approach is equivalent to looking at the microRNA strings from a holistic point of view. The other approach is to calculate exactly the probabilities of *n*-clusters of a particular state and fit them to the observed values.

#### 2.6.1. Cluster probabilities for large N Ising model

As mentioned earlier, we can verify the exactness of our model by checking the probabilities for high order clusters (11 and 00, 111 and 000 and so on). This is shown in Table 4. (Pu,St,Am) for 1’s and (Py,We,Ke) for 0’s. And the corresponding 2-D line plot is shown in Figure 5 in two different styles for (Pu, St, Am) and (Py, We, Ke). The *Pr*(*r*) values (*r* > 4) for Pu-Py class is more than the other classes.

**Table 4:** The calculated probability (*Pr*(*r*)) for different lengths strings, where *r* is the string of 1’s or 0’s.

|  | Pr(r) |  |  |  |  |  |  |  |
| --- | --- | --- | --- | --- | --- | --- | --- | --- |
| r= | 1 | 11 | 111 | 1111 | 11111 | 111111 | 1111111 | 11111111 |
| Pu | 0.509 | 0.292 | 0.173 | 0.104 | 0.062 | 0.038 | 0.025 | 0.016 |
| St | 0.522 | 0.251 | 0.122 | 0.061 | 0.032 | 0.019 | 0.012 | 0.008 |
| Am | 0.455 | 0.226 | 0.112 | 0.056 | 0.028 | 0.015 | 0.008 | 0.005 |
| r= | 0 | 00 | 000 | 0000 | 00000 | 000000 | 0000000 | 00000000 |
| Py | 0.491 | 0.269 | 0.157 | 0.097 | 0.061 | 0.039 | 0.027 | 0.019 |
| We | 0.478 | 0.197 | 0.091 | 0.043 | 0.019 | 0.008 | 0.004 | 0.002 |
| Ke | 0.545 | 0.314 | 0.179 | 0.102 | 0.059 | 0.014 | 0.020 | 0.012 |

In large N limit, the 1-cluster probability becomes (For the derivation and closed form expressions for higher-order clusters see App. 3.2),

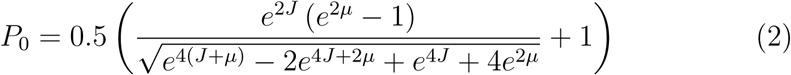

with

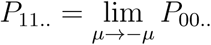

Eqn.s 2 and 16 (see App. 3.2) are solved with the observed probabilities for Pu-Py classification which give, *μ* = −0.0143414 and *J* = 0.113675.

Feeding these values in eqn.s 17 and 18 we find, *P*_000_ = 0.147573 (6% deviation from the observed value) and *P*_0000_ = 0.0809581 (16.5% deviation from the observed value).

However, doing the same thing with equations and probabilities for 1-clusters gives, *P*_111_ = 0.167168 (3.4% deviation from the observed value) and *P*_1111_ = 0.0957027 (8% deviation from the observed value).

#### 2.6.2. Cluster probabilities

In their paper [11] Ivanytskyi & Chelnokov have calculated the exact expression for average occupancy number of *distinct l*-clusters in Ising model in the absence of external field. The average occupancy number can be written as,

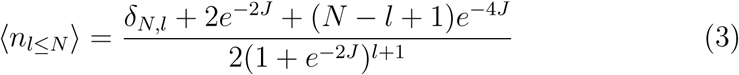

Now, the number of all distinct and non-distinct q-clusters, *ñ*_*q≤N*_, can be related to the number of all distinct clusters as,

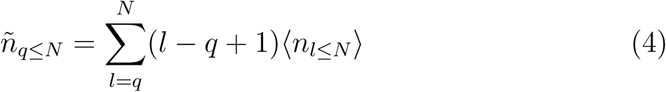

Solving equation 4 for *q* = 2 and *p*_00_ from Table 4 we get, *J* = 0.1728697. Using this *J* in equations 3 and 4 we predict *p*_000_ from this model to be 0.1714543, which deviates from the observation (Table 4) by 1.05% only.

#### 2.6.3. Variance from finite N Ising chain with PBC

In App. 3.1 we calculate a frequently used physical quantity, known as the isothermal susceptibility, for the Ising chain for finite N. This quantity can be argued to be identical to the variance of difference between the population of two classes (in case of Pu-Py classification, 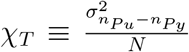). At *μ* → 0 limit this variance is given as,

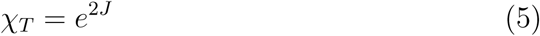

Solving for J with 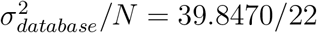 gives,

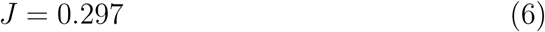

which is almost double the value obtained in subsection 2.6.2.

For the strong-weak H-bond grouping, 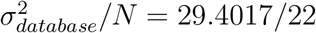 gives,

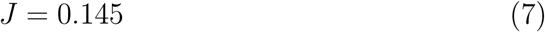

whereas, *J* = 0.175 for the amino-keto grouping.

## 3. Concluding Remarks

We observe that the Ising model with nearest neighbour interactions and chemical potential serves as a better descriptor of the miRNA data than the binomial model. We also observe that, within the scope of Ising model, the mean-field like quantities such as variance cannot lead us to a full description of the data. For a better description we may use a hybrid model which takes strong nearest-neighbour interactions into account with a background mean-field description.

At least for the Pu-Py classification, we observe that there is hardly any difference between the finite and large N Ising models in describing the miRNA data. So the finite number of elements in the string does not play an important role in our approximated description [12].

In all of our analysis we have mapped the four letter alphabet of nucleotide bases into a two letter binary alphabet consisting of zero and one. It is possible to keep the four letter alphabet and address the problem in all its generality. In that case the transfer matrix will have sixteen parameters. We may use an additive model based on the chemical classifications to lower the number of free parameters. We are currently working on this direction.

## Acknowledgments

The authors acknowledge late Prof. R. L. Brahmachary for his valuable suggestions.

## Appendix

### 3.1. App. - A: Nearest neighbor model

Here we solve the finite N Ising chain using transfer matrix method [12]. The transfer matrix in this problem is given by,

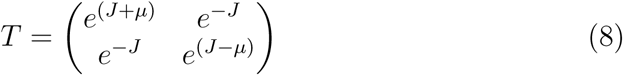

with the partition function, *Z* as,

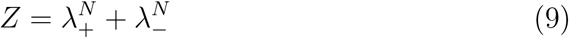

where

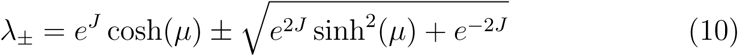

At *N* → ∞, the larger eigenvalue (*λ*_+_) dominates and the free energy density is given as,

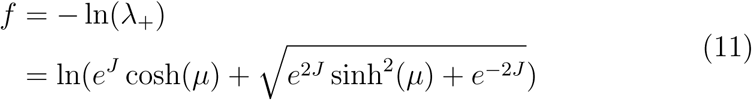

The isothermal susceptibility for this model is given by

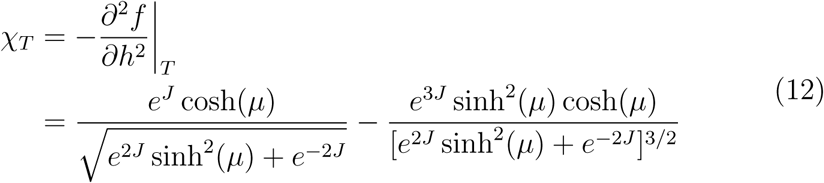

### 3.2. App. - B: Cluster Probabilities for large N

The cluster probabilities can be obtained using the transfer matrix method. For an n-cluster of 00.. the probability is obtained by using

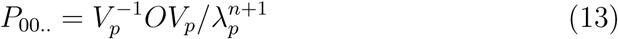

where *λ_p_* is the largest eigenvector of the transfer matrix, T, and *V_p_* is the corresponding eigenvector.

For *n* = 2,

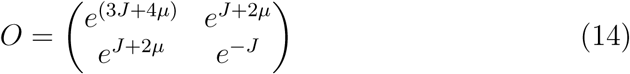

for *n* = 3,

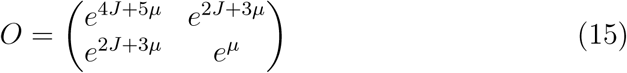

Using these we arrive at the probabilities for n-clusters,

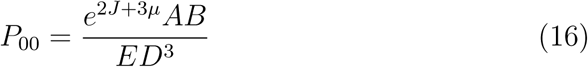

Where 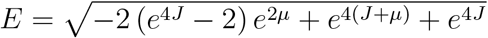
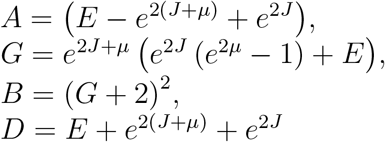

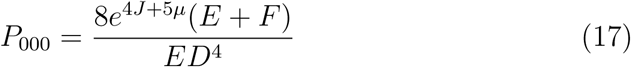

Where *F* = *e*^2*J*^ (*e*^2*μ*^ (*e^μ^* (*G* + 4) − 1) + 1),

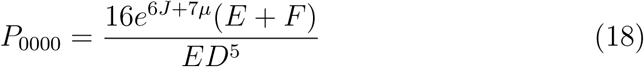

## References

[1] J. Li, Z. Zhang, mirna regulatory variation in human evolution, Trends in Genetics 29 (2) (2013) 116–124.

[2] S. Altman, Enzymatic cleavage of rna by rna, Bioscience reports 10 (4) (1990) 317–337.

[3] L. He, G. J. Hannon, Micrornas: small rnas with a big role in gene regulation, Nature reviews. Genetics 5 (8) (2004) 631.

[4] L. P. Lim, N. C. Lau, P. Garrett-Engele, A. Grimson, J. M. Schelter, J. Castle, D. P. Bartel, P. S. Linsley, J. M. Johnson, Microarray analysis shows that some micrornas downregulate large numbers of target mrnas, Nature 433 (7027) (2005) 769.

[5] L.-A. MacFarlane, P. R Murphy, Microrna: biogenesis, function and role in cancer, Current genomics 11 (7) (2010) 537–561.

[6] G. A. Calin, C. M. Croce, Microrna signatures in human cancers, Nature reviews cancer 6 (11) (2006) 857.

[7] J. K. Das, P. P. Choudhury, A. Chaudhuri, S. S. Hassan, P. Basu, Distribution of purines and pyrimidines over mirnas of human, gorilla and chimpanzee, bioRxiv (2017) 208405.

[8] L. Shi, H. Huang, Dna sequences analysis based on classifications of nucleotide bases, in: Affective Computing and Intelligent Interaction, Springer, 2012, pp. 379–384.

[9] Z. A. Shabarova, A. A. Bogdanov, Advanced organic chemistry of nucleic acids, John Wiley & Sons, 2008.

[10] A. Colliva, R. Pellegrini, A. Testori, M. Caselle, Ising-model description of long-range correlations in DNA sequences, pre 91 (5) (2015) 052703. arXiv:1409.0356, doi:10.1103/PhysRevE.91.052703.

[11] A. Ivanitskii, V. O. Chelnokov, On bimodal size distribution of spin clusters in the one dimensional ising model. URL https://arxiv.org/abs/1512.02676

[12] R. Pathria, P. Beale, Statistical Mechanics, Elsevier Science, 1996. URL https://books.google.co.in/books?id=PIk9sF9j2oUC

